# The Structure-Based Virtual Screening for Natural Compounds that Bind with the Activating Receptors of Natural Killer Cells

**DOI:** 10.1101/2020.06.19.160861

**Authors:** Adekunle Babajide Rowaiye, Solomon Oni, Ikemefuna Chijioke Uzochukwu, Alex Akpa, Charles Okechukwu Esimone

## Abstract

**Aim:** This study is aimed at prospecting for natural compounds that have strong binding affinity for the Activating Receptors of Natural Killer (NK) cells.

**Background:** NK cells are responsible for the immunosurveillance of tumor and virally- infected cells. The cytotoxic potentials of this unique population of immune cells are triggered by the activating receptors. Through ligand-binding, these receptors induce the tyrosine phosphorylation of adapter proteins through their Immunoreceptor Tyrosine–based Activation Motif ITAM sequences and this triggers direct cytotoxicity and the production of cytokines through different signal pathways.

**Objective:** To computationally predict the selectivity, specificity, and efficacy of natural compounds to be used as immunostimulatory agents for cancer treatment.

**Method:** In this study, 1,697 natural compounds were obtained from 82 edible tropical plants through data mining. The molecular docking simulations of these compounds were executed against 18 activating NK cells receptor targets using the Python Prescription 0.8. An arbitrary docking score ≥ −7.0 kcal/mol was chosen as cut off value. Further screening for oral bioavailability, promiscuity, molecular complexity and pharmacokinetic properties using the Swissadme and pkCSM webservers. The ligand similarity analysis and phylogenetic analysis of the receptors was carried out with the ChemMine and Clustal Omega webservers respectively. Binding site analyses and bioactivity prediction were also done with the Protein-Ligand Interaction Profiler and Molinspiration webservers respectively. Normal mode analyses were carried out with the CABS-flex 2.0 server.

**Result:** Seventeen bioactive and non-promiscuous lead compounds with good physicochemical and pharmacokinetic properties were identified.

**Conclusion:** Further tests are required to evaluate the efficacy of the lead compounds.

## 1.0 INTRODUCTION

Cancer is a group of diseases characterized by erratic cell growth which invade and spread into other parts of the body [1, 2]. They are caused by DNA damage and an ineffective DNA repair mechanism. According to a 2015 WHO report, Cancer is the second leading cause of death globally and there were 90.5 million incidences of cancer in 2015 which accounted for 8.8 million deaths [3].

Cancers are caused by a persistent damage to DNA which culminates into mutations of certain gene sequences in the human genome [4]. Expression of these mutant sequences lead to an autonomous and unregulated hyper-proliferation of cells; insufficient apoptosis; altered differentiation and metabolism; genomic instability and immortalization [5]. The abnormal proliferation of cells is due to alterations in the cell cycle replication mechanism due to nuclear and cytoplasm distortions. These changes include hyperchromatism, increased telomerase expression, prominent nucleoli, irregular chromatin distribution within nuclei, and increased size of nucleus, pleomorphism, and chromosomal translocations [6,7].

Available cancer therapies such as chemotherapy are non-selective as other normal rapidly dividing cells (including immune cells) are destroyed. Another major frustration faced by clinicians and researchers includes the evasive nature of cancer cells as they beat the immune system by their molecular ‘anonymity’. This is further complicated by their rapid multiplication, invasiveness and malignant abilities. Through intricate mechanisms, the rapidly dividing aberrant cells are able to evade the immune system, invade the surrounding tissue and enter into the lymph nodes and metastasize [8]. Therefore, the development of potential therapeutic agents must consider selectivity, specificity, and efficacy.

NK cells are responsible for immune-surveillance of tumor and virally infected cells. To unlock or lock the cytotoxic potentials of this unique population of immune cells are activating and inhibiting receptors respectively. The immunomodulatory potential of NK cells guarantees that the immune system does not fight against itself. Therefore, NK targeted therapies hold great promise in the treatment of cancers.

## CHAPTER 2: MATERIALS AND METHODS

### 2.1 Materials

The protein and ligand databases used were: Protein Databank, Uniprot, and PubChem. The webservers used were: pkCSM, Clustal Omega, ExPASy, Molinspiration, Protein-Ligand Interaction Profiler (PLIP), SWISS-MODEL, Swissadme, ChemMine, MolProbity, Chiron and CABS-flex 2.0. The softwares used were: Discovery studio 2017, Open babel, Pymol, and Python prescription (PyRx) 0.8.

### 2.2. Methods

#### 2.2.1 Identification of targets

The activating receptors of NK cells were identified by an extensive literature review. Validation of these molecular targets was also by empirical evidences provided by relevant research publications. The 3D crystallographic structures of these proteins were downloaded from the RCSB protein databank in the pdb format and visualized using the Pymol software [9]. The homology modeling of the proteins whose structures could not be obtained in the RCSB protein databank was executed using the SWISS-MODEL web-server [10]. The templates of closely related proteins were used for the modeling as seen in Table 1.

**Table 1:** Homology modeling of the proteins.

| S/N | Receptor name | Uniprot code | Template | % Similarity with template |
| --- | --- | --- | --- | --- |
| 1 | Killer cell immunoglobulin-like receptor 2DS1 | Q14954 | KIR3DL1 | 92.44 |
| 2 | Killer cell immunoglobulin-like receptor 2DS3 | Q14952 | KIR3DL1 | 88.36 |
| 3 | Killer cell immunoglobulin-like receptor 2DS5 | Q14953 | KIR2DL1 | 91.96 |
| 4 | NKG2-E type II integral membrane protein | Q07444 | NKG2A | 85.96 |

#### 2.2.2 Analysis and validation of protein structures

An all-atom structural validation and dihedral-angle diagnostics of the protein crystallography was conducted using the online server, MolProbity and the Ramanchandran plots were also obtained as seen in Table 2 [11].

**Table 2:** Ramanchandran Plot Analysis of Protein Structures.

| S/N | Receptor | Favoured Region (98%) | Allowed Region (>99.8%) | No of Outliers (%) | Outlier Residues |
| --- | --- | --- | --- | --- | --- |
| 1 | CD2 | 81.6% (84/103) | 96.1% (99/103) | 4 (0.039) | 8 GLU (-57.8, 91.6) |
|  |  |  |  |  | 27 SER (-167.0, -50.7) |
|  |  |  |  |  | 52 GLU (-177.9, 85.5) |
|  |  |  |  |  | 72 HIS (61.8, 110.6) |
| 2 | NCR2 | 94.3 (100/106) | 98.1(104/106) | 2 (0.019) | 59 TRP (-73.2, -140.7) |
|  |  |  |  |  | 60 THR (97.5, 67.6) |
| 3 | KIR2DS2 | 95.3% (182/191) | 97.9% (187/191) | 4(0.021) | 57 ASP (-66.3, 48.6) |
|  |  |  |  |  | 67 GLY (-29.0, 164.6) |
|  |  |  |  |  | 68 PRO (-33.4, 118.0) |
|  |  |  |  |  | 114 PRO (-61.5, -70.2) |
| 4 | NCR1 | 94.1% (175/186) | 98.9% (184/186) | 2 (0.011) | 100 TYR (60.2, -94.5) |
|  |  |  |  |  | 150 VAL (69.5, -34.2) |
| 5 | IL2Rα | 86.3% (101/117) | 96.6% (113/117) | 4 (0.034) | 22 GLU (-46.7, 99.6) |
|  |  |  |  |  | 112 ASN (-37.9, -169.8) |
|  |  |  |  |  | 116 GLU (150.2, -165.2) |
|  |  |  |  |  | 151 HIS (32.9, 78.8) |
| 6 | NKG2C | 82.0% (50/61) | 96.7% (59/61) | 2 (0.033) | 4 VAL (34.2, 31.5) |
|  |  |  |  |  | 33 LEU (46.3, 87.5) |
| 7 | KIR2DS4 | 90.2% (174/193) | 98.4% (190/193) | 3 (0.016) | 14 PRO (-64.1, -53.5) |
|  |  |  |  |  | 52 ILE (-91.4, 46.7) |
|  |  |  |  |  | 83 VAL (-118.9, -41.0) |
| 8 | NCR3 | 87.3% (96/110) | 100.0% (110/110) | 0 |  |
| 9 | IL2Rβ | 95.3% (183/192) | 100.0% (192/192) | 0 | |
| 10 | $\gamma c$ | 94.3% (181/192) | 100.0% (192/192) | 0 | |
| 11 | IL15Rα | 96.9% (63/65) | 100.0% (65/65) | 0 | |
| 12 | PILR | 94.1% (222/236) | 97.9% (231/236) | 5 (0.021) | A 2 LEU (-58.8, 30.2) |
|  |  |  |  |  | A 36 ASN (15.7, 74.4) |
|  |  |  |  |  | A 61 LYS (-25.5, -53.3) |
|  |  |  |  |  | B 36 ASN (18.0, 77.2) |
|  |  |  |  |  | B 61 LYS (-22.2, -59.2) |
| 13 | NKG2D | 93.1% (229/246) | 98.4% (242/246) | 4 (0.016) | A 116 GLU (-45.4, 83.0) |
|  |  |  |  |  | B 132 ALA (172.7, 134.5) |
|  |  |  |  |  | B 164 GLY (-53.6, 56.7) |
|  |  |  |  |  | B 176 PRO (-31.6, -74.6) |
| 14 | CD16 | 92.7% (140/151) | 100.0% (151/151) | 0 |  |
| 15 | KIR2DS1 | 87.3% (172/197) | 95.4% (188/197) | 9 (0.046) | 65 MET (-52.7, -74.3) |
|  |  |  |  |  | 78 ASP (-22.6, 112.8) |
|  |  |  |  |  | 88 SER (-48.2, 176.4) |
|  |  |  |  |  | 89 ARG (-22.7, 102.0) |
|  |  |  |  |  | 105 THR (-22.4, -53.6) |
|  |  |  |  |  | 107 SER (179.3, 55.2) |
|  |  |  |  |  | 135 PRO (-44.3, -50.9) |
|  |  |  |  |  | 163 GLU (-27.7, 101.6) |
|  |  |  |  |  | 166 ALA (-5.7, -67.1) |
| 16 | KIR2DS3 | 85.8% (169/197) | 96.4% (190/197) | 7(0.036) | 65 THR (-46.5, -75.6) |
|  |  |  |  |  | 78 ASP (-21.0, 113.3) |
|  |  |  |  |  | 89 ARG (-24.8, 104.3) |
|  |  |  |  |  | 107 SER (173.0, 54.7) |
|  |  |  |  |  | 135 PRO (-42.6, -53.0) |
|  |  |  |  |  | 163 GLU (-25.1, 99.9) |
|  |  |  |  |  | 166 ALA (-3.2, -70.1) |
| 17 | KIR2DS5 | 92.2% (178/193) | 98.4% (190/193) | 3 (0.016) | 105 THR (86.7, -37.8) |
|  |  |  |  |  | 188 ASP (92.4, -161.9) |
|  |  |  |  |  | 193 GLY (-56.8, -99.7) |
| 18 | NKG2E | 86.7% (98/113) | 99.1% (112/113) | 1(0.009) | 149 ASN (61.8, -81.9) |

#### 2.2.3 Preparation of protein targets for docking

In preparing the protein targets for molecular docking, all available water molecules, native ligands and unwanted chains were removed using the Pymol software [9]. Energy minimization of the protein targets to resolve steric clashes was done using online tool, *Chiron* as seen in Table 3 [12]. The PyRx software was used to convert the protein targets from pdb to pdbqt files [13].

**Table 3:** Energy Minimization of Protein Structures.

| S/<br>N | Protein | Structure | Total No. of<br>residues | Total No. of<br>contacts | Total No. of<br>clashes | Total VDW repulsion energy<br>(Kcal/mol) | Clash<br>ratio |
| --- | --- | --- | --- | --- | --- | --- | --- |
| 1 | CD2 | Initial | 105 | 1706 | 131 | 135.4 | 0.079 |
|  |  | Final | 105 | 1326 | 38 | 21.44 | 0.016 |
| 2 | NCR2 | Initial | 108 | 1539 | 55 | 39.48 | 0.026 |
|  |  | Final | 108 | 1497 | 37 | 26.14 | 0.017 |
| 3 | KIR2D<br>S2 | Initial | 193 | 2461 | 79 | 56.15 | 0.023 |
|  |  | Final | 193 | 2324 | 56 | 35.85 | 0.015 |
| 4 | NCR1 | Initial | 188 | 2714 | 84 | 53.82 | 0.02 |
|  |  | Final | 188 | 2714 | 84 | 53.82 | 0.02 |
| 5 | IL2Rα | Initial | 123 | 1520 | 65 | 52.26 | 0.034 |
|  |  | Final | 123 | 1378 | 39 | 24.39 | 0.018 |
| 6 | NKG2C | Initial | 63 | 788 | 51 | 54.34 | 0.07 |
|  |  | Final | 63 | 701 | 14 | 11.35 | 0.016 |
| 7 | KIR2D<br>S4 | Initial | 195 | 2637 | 103 | 85.66 | 0.032 |
|  |  | Final | 195 | 2398 | 69 | 42.26 | 0.018 |
| 8 | NCR3 | Initial | 112 | 1450 | 53 | 38.42 | 0.026 |
|  |  | Final | 112 | 1394 | 41 | 22.95 | 0.016 |
| 9 | IL2Rβ | Initial | | | | | |
|  |  | Final |  |  |  |  |  |
| 10 | $\gamma c$ | Initial | 193 | 2732 | 67 | 39.86 | 0.015 |
|  |  | Final | 193 | 2732 | 67 | 39.86 | 0.015 |
| 11 | IL15Rα | Initial | 67 | 847 | 25 | 17 | 0.02 |
|  |  | Final | 67 | 843 | 25 | 16.22 | 0.019 |
| 12 | PILR | Initial | 240 | 3324 | 101 | 66.67 | 0.02 |
|  |  | Final | 240 | 3402 | 93 | 57.85 | 0.017 |
| 13 | NKG2D | Initial | 250 | 4424 | 121 | 84.66 | 0.02 |
|  |  | Final | 250 | 4041 | 115 | 71.66 | 0.018 |
| 14 | CD16 | Initial | 157 | 2058 | 97 | 79 | 0.038 |
|  |  | Final | 157 | 2002 | 54 | 35.25 | 0.018 |
| 15 | KIR2D<br>S1 | Initial | 199 | 2507 | 65 | 37.54 | 0.015 |
|  |  | Final | 199 | 2507 | 65 | 37.54 | 0.015 |
| 16 | KIR2D<br>S3 | Initial | 199 | 2452 | 73 | 39.33 | 0.016 |
|  |  | Final | 199 | 2452 | 73 | 39.33 | 0.016 |
| 17 | KIR2D<br>S5 | Initial | 195 | 2281 | 55 | 28.65 | 0.013 |
|  |  | Final | 195 | 2281 | 55 | 28.65 | 0.013 |
| 18 | NKG2E | Initial | 115 | 1459 | 27 | 16.99 | 0.012 |
|  |  | Final | 115 | 1459 | 27 | 16.99 | 0.012 |

#### 2.2.4 Building of library of natural bioactive compounds

A library of 1,697 compounds was built from an extensive data mining from the literature review of 79 plants (See supplementary data) predominantly found in Nigeria and tropical Africa. The 3D structures of these natural compounds were downloaded from the PubChem chemical database in their SDF format [14]. The properties of these compounds such as molecular weight, canonical SMILES, number of heavy atoms, hydrogen bond donors, hydrogen bond acceptors, Log P and topological polar surface area were obtained from Pubchem [14].

#### 2.2.5 Preparation for docking

Prior to docking, 1697 natural compounds were screened for bioavailability using the Lipinski and Veber rules. As stated by Lipinski, the drug-like properties include a MW ≥ 500, Hydrogen Bond Donor ≥ 10, Hydrogen Bond Acceptor ≥ 5 and a Log P value ≥ 5. Further screening was done for cellular permeability using the Veber’s rule. Only compounds of Topological Polar Surface Area (TPSA) values of ≥ 140 and number of rotatable bonds ≥ 10 were successful [28, 29].

The docking protocol was validated by using a structure from the Protein Data Bank. The molecule which is the Adhesion Domain of Human CD2 (PDB ID: 1GYA) was downloaded in pdb format and separated from N-Glycan which is the native ligand. The separated molecules were docked together using PyRx 0.8. The docked result was superimposed on the pure protein structure and compared with the original 1GYA structure found in the data bank (Figure 1).

**Figure 1:**
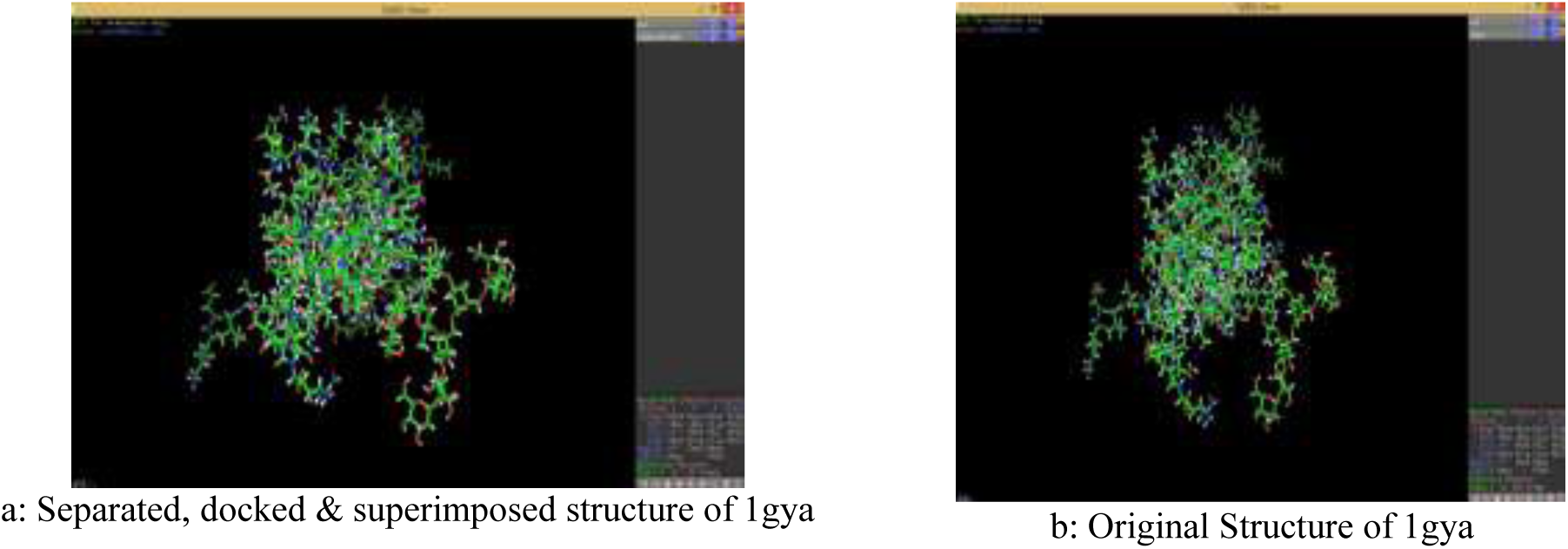
Separated, docked and superimposed structure of 1gya as compared to the original structure.

Ligands were uploaded unto PyRx 0.8 through the Open babel plug-in. For stable conformation, the conjugate gradient descent was used as optimization algorithm. The Universal Force Field (UFF) was used as the energy minimization parameter.

The Spatial Data File (SDF) formats of all ligands were converted to the pdbqt format in readiness for docking. The grids were maximized to cover the entire binding site of the ligand. Molecular docking of ligands against protein targets was executed through AutoDock Vina plug-in of the PyRx software. Based on the scoring function, the best fits were obtained and saved in excel files.

#### 2.2.6 Screening for potency

The first stage of the screening was for drug potency. Molecular docking was used as the first step in the virtual screening process and the docking scores were used as empirical predictors of the strength of the intermolecular interactions between the receptors and the ligands (See supplementary data).

A uniform docking scoring cutoff of −7.0 kcal/mol was used to serve as a general border line for the binding energies obtained between the receptors and the ligands. Because drug potency is an aggregate of the binding affinity and the efficacy, further screening for efficacy was executed by imploring the use of three Ligand Efficiency Metrics (LEM) which are the Ligand Efficiency (LE), ligand-efficiency-dependent lipophilicity, (LELP) and Ligand-lipophilicity efficiency (LLE). The LE was calculated as the binding energy divided by the number of heavy atoms; the LELP is the Log P value of the ligand divided by the LE; and the LLE is the binding energy minus the log P. The cut offs are ≥ 0.3 for LE; −10 to 10 for LELP; and ≥ 5.0 for LLE (See supplementary data for results).

#### 2.2.7 Further screening for Oral Bioavailability, Promiscuity and pharmacokinetic properties

After the initial screening for drug likeness using the Lipinski and Veber rules, the natural compounds were screened for saturation and promiscuity using the SWISSADME webserver [15]. Using the canonical SMILES, a Quantitative-Structural Activity Relationship (QSAR) based prediction of the Absorption, Distribution, Metabolism, Excretion and Toxicity (ADMET) properties of the selected compounds was executed using the pkCSM and this was used for further screening. (See supplementary data for results).

#### 2.2.8: Prediction of Bioactivity

Using the Molinspiration webserver, the bioactivity of the compounds was predicted as seen in Table 10.

**Table 4:** Lead compounds’ compliance with Lipinski & Veber rules.

|  | Pubchem ID | MW(g/mol) | Log P | HBD | HBA | TPSA (A<sup>2</sup>) | No of Rotatable bonds |
| --- | --- | --- | --- | --- | --- | --- | --- |
| 4'-Methyl-epigallocatechin | 176920 | 320.29 | 0.3 | 5 | 7 | 120 | 2 |
| Andrographis Extract | 6436016 | 350.4 | 2.2 | 3 | 5 | 87 | 3 |
| Betulalbuside A | 14484636 | 332.39 | -0.3 | 5 | 7 | 120 | 8 |
| Bisabolone Oxide A | 91700388 | 236.35 | 2.5 | 0 | 2 | 26.3 | 1 |
| Carveol | 7438 | 152.23 | 2.1 | 1 | 1 | 20.2 | 1 |
| cis-Carvotanacetol | 12233170 | 154.25 | 2.1 | 1 | 1 | 20.2 | 1 |
| Eugenyl Glucoside | 3084296 | 326.34 | 0 | 4 | 7 | 109 | 6 |
| Gibberellin A17 | 5460657 | 378.4 | 0.8 | 4 | 7 | 132 | 3 |
| Gibberellin A19 | 5460209 | 362.4 | 0.7 | 3 | 6 | 112 | 3 |
| Gibberellin A20 | 5280481 | 332.4 | 1.2 | 2 | 5 | 83.8 | 1 |
| Gibberellin A29 | 5460028 | 348.4 | 0.2 | 3 | 6 | 104 | 1 |
| Gibberellin A44 | 5460372 | 346.4 | 1.6 | 2 | 5 | 83.8 | 1 |
| Gibberellin A51 | 443458 | 332.4 | 1.7 | 2 | 5 | 83.8 | 1 |
| Gibberellin A53 | 440914 | 348.4 | 2.2 | 3 | 5 | 94.8 | 2 |
| Monocrotalline | 9415 | 325.36 | -0.7 | 2 | 7 | 96.3 | 0 |
| Phellandrenol | 76373091 | 152.23 | 2 | 1 | 1 | 20.2 | 2 |
| Shikimic Acid | 8742 | 174.15 | -1.7 | 4 | 5 | 98 | 1 |

**Table 5:** Summary of screening results.

|  |  |
| --- | --- |
| Total Library of compounds | 1697 |
| Bioavailability screening | 1048 |
| Docking results cut off | 377 |
| Ligand Efficiency Metrics screening | 192 |
| Promiscuity and Pharmacokinetics screening | 69 |
| Bioactivity screening | 17 |

**Table 6:**
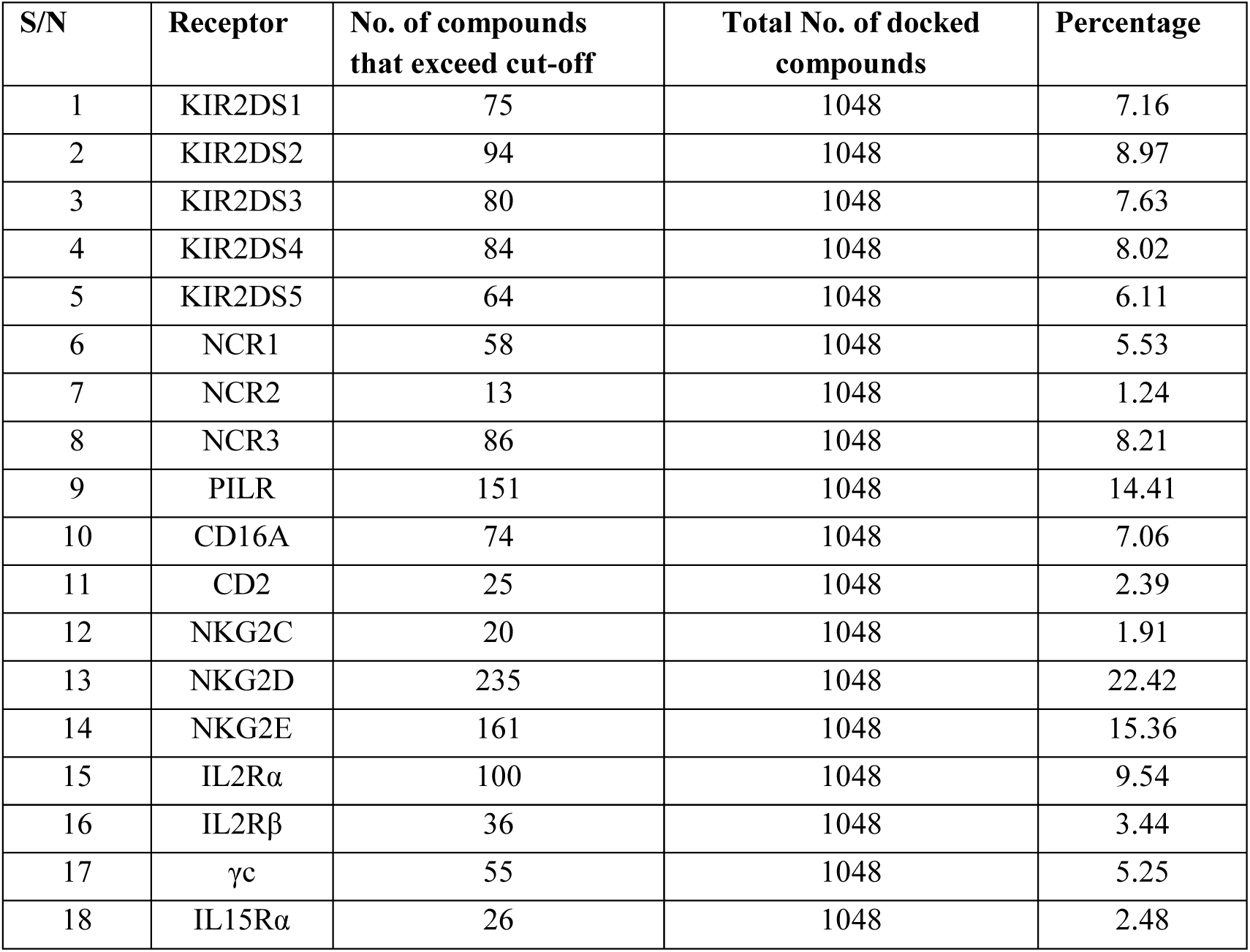
Summary of distributions and frequencies of receptor - ligand dockings (≤ −7.0kcal/mol)

**Table 7:** Absorption and Distribution profile of lead compounds.

| Ligand | H<sub>2</sub> O<br>Solubility | Caco2<br>perm. | HIA | Skin<br>Perm | P-gp<br>sub. | P-gp<br>I Inb. | P-gp<br>II Inb. | VDss | Fraction<br>unbound | BBB<br>perm | CNS<br>perm |
| --- | --- | --- | --- | --- | --- | --- | --- | --- | --- | --- | --- |
| 4'-Methyl-epigallocatechin | -3.09 | -0.12 | 60.7<br>3 | -2.74 | Yes | No | No | 1.64 | 0.26 | -0.93 | -3.27 |
| Andrographis Extract | -3.49 | 1.07 | 95.3<br>6 | -3.79 | No | No | No | -0.29 | 0.28 | -0.6 | -2.69 |
| Betulalbuside A | -1.94 | -0.14 | 43.9<br>6 | -3.03 | No | No | No | -0.26 | 0.65 | -1.01 | -3.65 |
| Bisabolone Oxide A | -3.63 | 1.62 | 96.4<br>7 | -2.52 | No | No | No | 0.34 | 0.43 | 0.55 | -3.05 |
| Carveol | -1.78 | 1.4 | 95.1<br>8 | -2.08 | No | No | No | 0.17 | 0.55 | 0.56 | -2.58 |
| cis-Carvotanacetol | -2.15 | 1.37 | 95.1<br>7 | -1.93 | No | No | No | 0.13 | 0.47 | 0.58 | -2.12 |
| Eugenyl Glucoside | -1.89 | 0.58 | 45.7<br>1 | -2.87 | Yes | No | No | -0.38 | 0.41 | -0.99 | -3.73 |
| Gibberellin A17 | -2.89 | 0.82 | 33.0<br>3 | -2.74 | No | No | No | -0.89 | 0.42 | -0.77 | -3.31 |
| Gibberellin A19 | -2.81 | 0.96 | 47.5 | -2.74 | No | No | No | -1.6 | 0.42 | -0.68 | -3.16 |
| Gibberellin A20 | -2.64 | 1.19 | 98.9<br>1 | -2.74 | No | No | No | -0.83 | 0.42 | -0.21 | -3 |
| Gibberellin A29 | -2.66 | 0.69 | 71.4<br>2 | -2.74 | No | No | No | -0.82 | 0.47 | -0.6 | -3.11 |
| Gibberellin A44 | -2.84 | 1.18 | 99.3 | -2.74 | No | No | No | -1.11 | 0.3 | -0.18 | -2.29 |
| Gibberellin A51 | -2.73 | 1.14 | 100 | -2.74 | No | No | No | -0.97 | 0.29 | -0.09 | -2.41 |
| Gibberellin A53 | -2.82 | 0.93 | 53.2<br>7 | -2.74 | No | No | No | -1.6 | 0.36 | -0.56 | -2.31 |
| Monocrotaline | -3.06 | 0.51 | 64.7<br>6 | -3.01 | Yes | No | No | 0.47 | 0.74 | -0.61 | -3.1 |
| Phellandrenol | -1.96 | 1.49 | 94.9<br>9 | -2.33 | Yes | No | No | 0.22 | 0.56 | 0.55 | -2.69 |
| Shikimic Acid | -0.52 | -0.23 | 46.6<br>8 | -2.74 | No | No | No | -0.62 | 0.8 | -0.68 | -3.58 |

HIA= Human Intestinal Absorption. Skin Perm = Skin Permeability. P-gp = Plasma glycoprotein. VDSS= Volume of Distribution steady State. BBB= Blood Brain Barrier

**Table 8:** Metabolism, Excretion, and Saturation and Agglutination profile of lead compounds.

| Ligand | CYP2D<br>6 sub | CYP3A<br>4 sub | CYP1A<br>2 inh | CYP2C<br>19 inh | CYP2C<br>9 inh | CYP2D<br>6 inh | CYP3A<br>4 inh | Total<br>Clearan<br>ce | Renal<br>OCT<br>2 sub | Fractio<br>n Csp3 | PAIN<br>S<br>#alert<br>s |
| --- | --- | --- | --- | --- | --- | --- | --- | --- | --- | --- | --- |
| 4'-Methyl-epigallocatechin | No | No | No | No | No | No | No | 0.35 | No | 0.25 | 0 |
| Andrographis Extr<br>act | NO | Yes | No | No | No | No | No | 1.18 | No | 0.75 | 0 |
| Betulalbuside A | No | No | No | No | No | No | No | 1.69 | No | 0.75 | 0 |
| Bisabolone Oxide<br>A | No | No | No | No | No | No | No | 1.13 | No | 0.8 | 0 |
| Carveol | No | No | No | No | No | No | No | 0.23 | No | 0.6 | 0 |
| cis-Carvotanacetol | No | No | No | No | No | No | No | 0.19 | No | 0.8 | 0 |
| Eugenyl Glucosid<br>e | No | No | No | No | No | No | No | 0.26 | No | 0.5 | 0 |
| Gibberellin A17 | No | No | No | No | No | No | No | 0.39 | No | 0.75 | 0 |
| Gibberellin A19 | NO | Yes | No | No | No | No | No | 0.47 | No | 0.75 | 0 |
| Gibberellin A20 | No | Yes | No | No | No | No | No | 0.42 | No | 0.79 | 0 |
| Gibberellin A29 | No | Yes | No | No | No | No | No | 0.42 | No | 0.79 | 0 |
| Gibberellin A44 | No | Yes | No | No | No | No | No | 0.36 | No | 0.8 | 0 |
| Gibberellin A51 | No | Yes | No | No | No | No | No | 0.42 | No | 0.79 | 0 |
| Gibberellin A53 | No | Yes | No | No | No | No | No | 0.43 | No | 0.8 | 0 |
| Monocrotaline | No | No | No | No | No | No | No | 0.73 | No | 0.75 | 0 |
| Phellandrenol | No | No | No | No | No | No | No | 0.29 | No | 0.6 | 0 |
| Shikimic Acid | No | No | No | No | No | No | No | 0.69 | No | 0.57 | 0 |

Renal OCT2 = Renal Organic Cation transporter

**Table 9:** Toxicity profile of lead compounds.

| Ligand | AME<br>S<br>toxicit<br>y | Max.<br>tolerat<br>ed dose | hERG<br>G I<br>inh | hERG<br>G II<br>inh | Oral<br>Rat<br>Acute<br>Toxicity<br>(LD50<br>) | Oral<br>Rat<br>Chronic<br>Toxicity<br>(LOAE<br>L) | Hepatotoxic<br>ity | Skin<br>Sensitisation | T.Pyriformis toxicity | Minnow<br>toxicity |
| --- | --- | --- | --- | --- | --- | --- | --- | --- | --- | --- |
| 4'-Methyl-epigallocatechin | No | 0.37 | No | NO | 2.29 | 2.93 | No | No | 0.3 | 3.75 |
| Andrographis Extract | No | 0.13 | No | No | 2.16 | 1 | No | No | 0.49 | 1.37 |
| Betulalbuside A | No | 1.37 | No | NO | 1.71 | 3.2 | No | No | 0.29 | 3.66 |
| Bisabolone Oxide A | No | 0.35 | No | No | 1.99 | 1.86 | No | Yes | 0.73 | 1.07 |
| Carveol | No | 0.84 | No | No | 1.96 | 1.89 | No | Yes | 0.2 | 1.67 |
| cis-Carvotanacetol | No | 0.82 | No | No | 1.98 | 1.99 | No | Yes | 0.32 | 1.36 |
| Eugenyl Glucoside | No | 0.86 | No | No | 1.95 | 3.46 | Yes | No | 0.29 | 3.8 |
| Gibberellin A17 | No | 0.44 | No | No | 2.48 | 2.7 | No | No | 0.29 | 3.1 |
| Gibberellin A19 | No | 0.4 | No | No | 2.21 | 2.28 | Yes | No | 0.29 | 2.49 |
| Gibberellin A20 | No | 0.37 | No | No | 2.05 | 2.14 | No | No | 0.29 | 1.96 |
| Gibberellin A29 | No | 0.26 | No | NO | 2.1 | 2.5 | No | No | 0.29 | 2.69 |
| Gibberellin A44 | No | 0.15 | No | NO | 2.06 | 1.96 | No | No | 0.29 | 1.76 |
| Gibberellin A51 | No | -0.14 | No | NO | 2.1 | 2.36 | Yes | No | 0.29 | 1.16 |
| Gibberellin A53 | No | 0.36 | No | No | 2.2 | 2.17 | Yes | No | 0.29 | 1.76 |
| Monocrotaline | No | 0.42 | No | No | 2.4 | 1.99 | No | No | 0.29 | 3.88 |
| Phellandrenol | No | 0.87 | No | No | 1.83 | 1.81 | No | Yes | 0.09 | 1.72 |
| Shikimic Acid | No | 0.99 | No | No | 1.16 | 2.96 | No | No | 0.26 | 4.05 |

hERG = human Ether-a-go-go-related Gene.

**Table 10:** Bioactivity profile of front-runner compounds.

| S/N | Ligand | GPC<br>R<br>ligan<br>d | Ion<br>channel<br>mod. | *Kinase*<br>*Inh.* | Nuclear<br>Receptor<br>Ligand | Protease<br>Inh. | Enzyme<br>Inh | No of Recep.<br>Targets ($\leq -7.0$ kcal/mol) |
| --- | --- | --- | --- | --- | --- | --- | --- | --- |
| 1 | Andrographis Extract | 0.32 | 0.17 | -0.01 | 0.94 | 0.26 | 0.81 | 16 |
| 2 | Gibberellin A53 | 0.39 | 0.17 | -0.35 | 0.76 | 0.18 | 0.42 | 6 |
| 3 | Gibberellin A19 | 0.32 | 0.10 | -0.30 | 0.69 | 0.30 | 0.43 | 15 |
| 4 | Gibberellin A51 | 0.17 | 0.21 | -0.31 | 0.67 | 0.16 | 0.38 | 14 |
| 5 | Gibberellin A44 | 0.34 | 0.16 | -0.21 | 0.66 | 0.19 | 0.36 | 16 |
| 6 | Gibberellin A17 | 0.36 | 0.11 | -0.25 | 0.63 | 0.18 | 0.33 | 17 |
| 7 | Gibberellin A29 | 0.24 | 0.20 | -0.24 | 0.60 | 0.19 | 0.42 | 15 |
| 8 | Gibberellin A20 | 0.22 | 0.23 | -0.21 | 0.49 | 0.09 | 0.30 | 7 |
| 9 | 4'-Methyl-epigallocatechin | 0.37 | 0.07 | 0.11 | 0.48 | 0.23 | 0.39 | 17 |
| 10 | Monocrotaline | 0.36 | 0.38 | -0.05 | 0.47 | 0.50 | 0.28 | 7 |
| 11 | Betulalbuside A | 0.27 | 0.35 | -0.05 | 0.38 | 0.22 | 0.73 | 1 |
| 12 | Carveol | -0.55 | 0.14 | -1.40 | 0.25 | -0.89 | 0.23 | 1 |
| 13 | Bisabolone Oxide A | -0.11 | 0.10 | -0.97 | 0.24 | -0.35 | 0.56 | 1 |
| 14 | Phellandrenol | -0.75 | -0.34 | -1.07 | 0.12 | -1.14 | 0.23 | 1 |
| 15 | Eugenyl Glucoside | 0.05 | -0.03 | -0.21 | 0.02 | -0.11 | 0.32 | 2 |
| 16 | Shikimic Acid | -0.38 | 0.22 | -1.13 | 0.01 | -0.37 | 0.65 | 1 |
| 17 | cis-Carvotanacetol | -0.50 | 0.09 | -1.09 | 0.01 | -0.62 | 0.18 | 1 |

#### 2.2.9: Specificity/promiscuity analyses

After the initial screenings, the comparative binding affinity analysis of all the protein-ligand interactions was done to check for specific and promiscuous binding (Table 11).

**Table 11:**
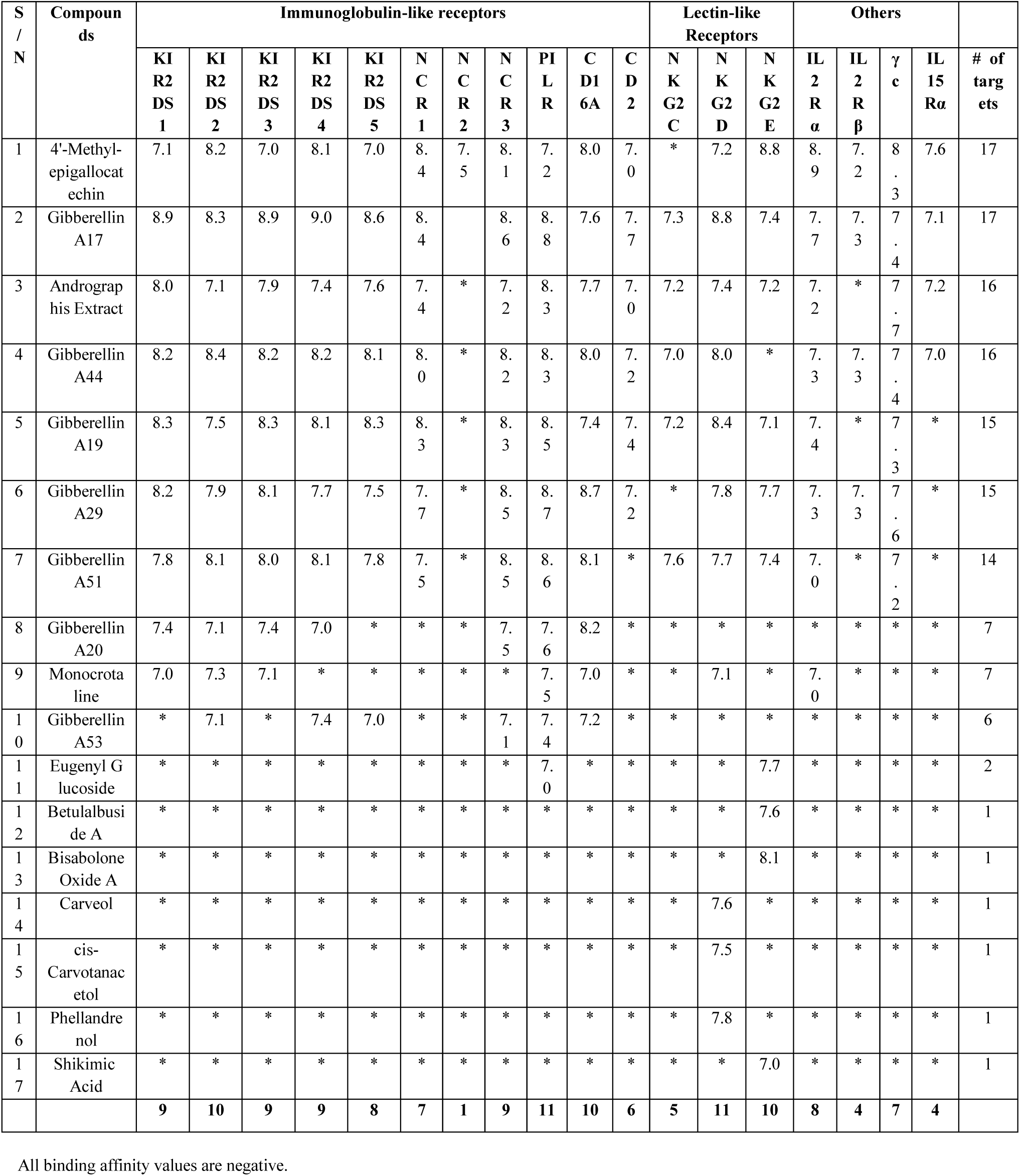
Binding affinities of front runner compounds (post-screening) with cut off value of ≤ −7.0 kcal/mol.

#### 2.2.10 Structural similarity analyses

The similarity analyses of all the screened ligands were done using the *ChemMine* webserver as shown in Table 12 [16]. A structural analysis of the protein targets was done through a pairwise Percent Identity Matrix. The results are seen in Table 13. A multiple sequence alignment of the amino acid residues of the extracellular domain of all the receptor targets and subsequently the phylogenetic analysis was done using the Clustal Omega webserver [17]. The results are shown in Figure 2.

**Table 12:**
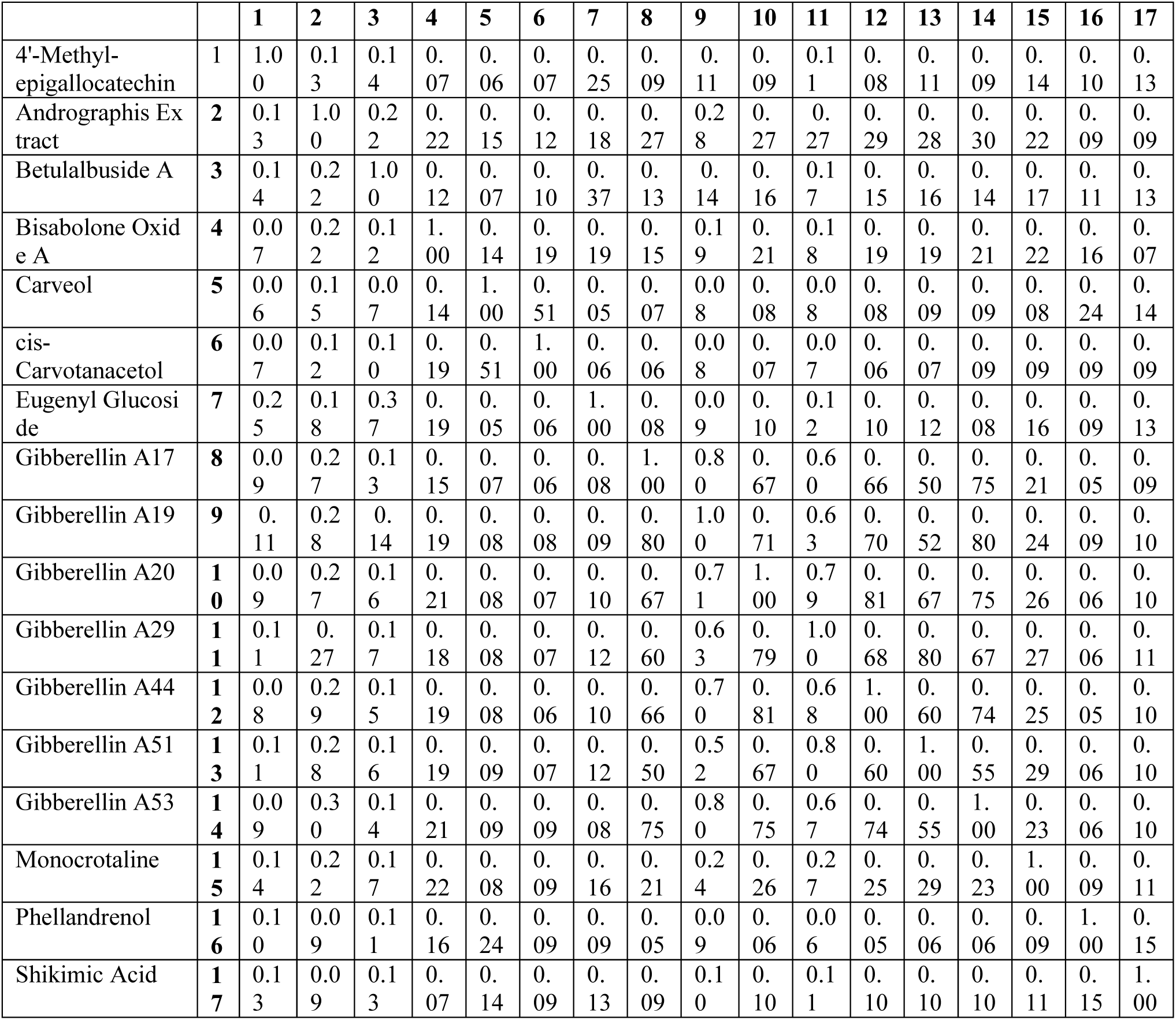
Pairwise Ligand Similarity Analysis of active compounds Using Tanimoto Coefficient.

**Table 13:** Percent Identity Matrix of Protein Targets.

|  |  | 1 | 2 | 3 | 4 | 5 | 6 | 7 | 8 | 9 | 10 | 11 | 12 | 13 | 14 | 15 | 16 | 17 | 18 |
| --- | --- | --- | --- | --- | --- | --- | --- | --- | --- | --- | --- | --- | --- | --- | --- | --- | --- | --- | --- |
| **NCR2** | **1** | 10<br>0 | 25.<br>12 | 15.<br>56 | 15.<br>38 | 21.<br>69 | 12.<br>79 | 10.<br>26 | 15.<br>19 | 17.<br>24 | 21.<br>57 | 10.<br>85 | 17.<br>69 | 14.<br>29 | 12.<br>82 | 12.<br>82 | 12.<br>82 | 11.<br>97 | 11.<br>97 |
| **IL15R**<br>**α** | **2** | 25.<br>12 | 10<br>0 | 15.<br>2 | 14.<br>09 | 17.<br>13 | 14.<br>52 | 19.<br>64 | 21.<br>05 | 10.<br>59 | 14.<br>65 | 10.<br>45 | 13.<br>48 | 15.<br>13 | 13.<br>73 | 13.<br>73 | 13.<br>73 | 14.<br>38 | 13.<br>73 |
| **IL2Rα** | **3** | 15.<br>56 | 15.<br>2 | 10<br>0 | 21.<br>08 | 17.<br>91 | 11.<br>29 | 11.<br>27 | 11.<br>27 | 13.<br>79 | 18.<br>09 | 8.4<br>6 | 9.7<br>6 | 6.5<br>7 | 8.4<br>5 | 7.7<br>5 | 9.1<br>5 | 8.4<br>5 | 8.4<br>5 |
| **NCR3** | **4** | 15.<br>38 | 14.<br>09 | 21.<br>08 | 10<br>0 | 26.<br>23 | 12.<br>16 | 12.<br>5 | 12.<br>33 | 13.<br>57 | 12.<br>86 | 9.6<br>2 | 11.<br>58 | 10.<br>28 | 12.<br>38 | 13.<br>33 | 13.<br>33 | 15.<br>24 | 13.<br>33 |
| **PILRB** | **5** | 21.<br>69 | 17.<br>13 | 17.<br>91 | 26.<br>23 | 10<br>0 | 16.<br>95 | 12.<br>07 | 12.<br>07 | 13.<br>53 | 12.<br>5 | 10.<br>16 | 11.<br>81 | 12.<br>59 | 10.<br>42 | 13.<br>19 | 11.<br>11 | 11.<br>81 | 11.<br>81 |
| **NKG2**<br>**D** | **6** | 12.<br>79 | 14.<br>52 | 11.<br>29 | 12.<br>16 | 16.<br>95 | 10<br>0 | 22.<br>06 | 22.<br>33 | 15 | 15.<br>12 | 13.<br>21 | 10.<br>58 | 13.<br>6 | 11.<br>11 | 10.<br>1 | 11.<br>11 | 11.<br>11 | 9.0<br>9 |
| **NKG2**<br>**C** | **7** | 10.<br>26 | 19.<br>64 | 11.<br>27 | 12.<br>5 | 12.<br>07 | 22.<br>06 | 10<br>0 | 90.<br>04 | 9.4<br>9 | 21.<br>55 | 8.2<br>6 | 11.<br>21 | 16.<br>28 | 17.<br>92 | 17.<br>92 | 15.<br>09 | 19.<br>81 | 17.<br>92 |
| **NKG2**<br>**E** | **8** | 15.<br>19 | 21.<br>05 | 11.<br>27 | 12.<br>33 | 12.<br>07 | 22.<br>33 | 90.<br>04 | 100 | 8.7<br>6 | 21.<br>16 | 6.3<br>6 | 11.<br>21 | 14.<br>73 | 16.<br>98 | 16.<br>98 | 14.<br>15 | 18.<br>87 | 16.<br>98 |
| **Comm**<br>**on γc** | **9** | 17.<br>24 | 10.<br>59 | 13.<br>79 | 13.<br>57 | 13.<br>53 | 15 | 9.4<br>9 | 8.7<br>6 | 100 | 22.<br>22 | 8.4<br>3 | 11.<br>45 | 9.8<br>5 | 14.<br>75 | 13.<br>11 | 13.<br>66 | 14.<br>21 | 13.<br>11 |
| **IL2Rβ** | **10** | 21.<br>57 | 14.<br>65 | 18.<br>09 | 12.<br>86 | 12.<br>5 | 15.<br>12 | 21.<br>55 | 21.<br>16 | 22.<br>22 | 10<br>0 | 17.<br>52 | 18.<br>35 | 13.<br>28 | 13.<br>27 | 14.<br>29 | 14.<br>29 | 13.<br>27 | 13.<br>78 |
| **CD2** | **11** | 10.<br>85 | 10.<br>45 | 8.4<br>6 | 9.6<br>2 | 10.<br>16 | 13.<br>21 | 8.2<br>6 | 6.3<br>6 | 8.4<br>3 | 17.<br>52 | 10<br>0 | 18.<br>63 | 19.<br>07 | 19.<br>68 | 20.<br>74 | 20.<br>21 | 18.<br>62 | 19.<br>68 |
| **CD16A** | **12** | 17.<br>69 | 13.<br>48 | 9.7<br>6 | 11.<br>58 | 11.<br>81 | 10.<br>58 | 11.<br>21 | 11.<br>21 | 11.<br>45 | 18.<br>35 | 18.<br>63 | 10<br>0 | 21.<br>2 | 19.<br>62 | 19.<br>14 | 20.<br>1 | 20.<br>1 | 21.<br>53 |
| **NCR1** | **13** | 14.<br>29 | 15.<br>13 | 6.5<br>7 | 10.<br>28 | 12.<br>59 | 13.<br>6 | 16.<br>28 | 14.<br>73 | 9.8<br>5 | 13.<br>28 | 19.<br>07 | 21.<br>2 | 10<br>0 | 34.<br>76 | 34.<br>76 | 34.<br>33 | 33.<br>91 | 33.<br>48 |
| **KIR2D**<br>**S4** | **14** | 12.<br>82 | 13.<br>73 | 8.4<br>5 | 12.<br>38 | 10.<br>42 | 11.<br>11 | 17.<br>92 | 16.<br>98 | 14.<br>75 | 13.<br>27 | 19.<br>68 | 19.<br>62 | 34.<br>76 | 10<br>0 | 89.<br>39 | 90.<br>2 | 86.<br>53 | 86.<br>94 |
| **KIR2D**<br>**S1** | **15** | 12.<br>82 | 13.<br>73 | 7.7<br>5 | 13.<br>33 | 13.<br>19 | 10.<br>1 | 17.<br>92 | 16.<br>98 | 13.<br>11 | 14.<br>29 | 20.<br>74 | 19.<br>14 | 34.<br>76 | 89.<br>39 | 10<br>0 | 92.<br>65 | 91.<br>43 | 92.<br>24 |
| **KIR2D**<br>**S2** | **16** | 12.<br>82 | 13.<br>73 | 9.1<br>5 | 13.<br>33 | 11.<br>11 | 11.<br>11 | 15.<br>09 | 14.<br>15 | 13.<br>66 | 14.<br>29 | 20.<br>21 | 20.<br>1 | 34.<br>33 | 90.<br>2 | 92.<br>65 | 10<br>0 | 91.<br>84 | 91.<br>43 |
| **KIR2D**<br>**S3** | **17** | 11.<br>97 | 14.<br>38 | 8.4<br>5 | 15.<br>24 | 11.<br>81 | 11.<br>11 | 19.<br>81 | 18.<br>87 | 14.<br>21 | 13.<br>27 | 18.<br>62 | 20.<br>1 | 33.<br>91 | 86.<br>53 | 91.<br>43 | 91.<br>84 | 10<br>0 | 92.<br>65 |
| **KIR2D**<br>**S5** | **18** | 11.<br>97 | 13.<br>73 | 8.4<br>5 | 13.<br>33 | 11.<br>81 | 9.0<br>9 | 17.<br>92 | 16.<br>98 | 13.<br>11 | 13.<br>78 | 19.<br>68 | 21.<br>53 | 33.<br>48 | 86.<br>94 | 92.<br>24 | 91.<br>43 | 92.<br>65 | 10<br>0 |

#### 2.2.11: Binding Site analyses

The poses of the selected ligands as they interact with the receptors during docking were saved on PyRx and viewed on PyMol. The protein structures were superimposed on PyMol and saved in the pdb format. The structures were uploaded into the Protein-Ligand Interaction Profiler (PLIP) webserver for the analysis of their binding sites [18]. The summary of all the protein-ligand interactions are shown in supplementary data.

#### 2.2.12: Normal Mode Analysis

The Root Mean Square Fluctuation (RMSF) plots of the amino acid residues of native and mutant (after binding with ligand) proteins were obtained using the CABS-flex 2.0 webserver (Table 14) [19].

**Table 14:**
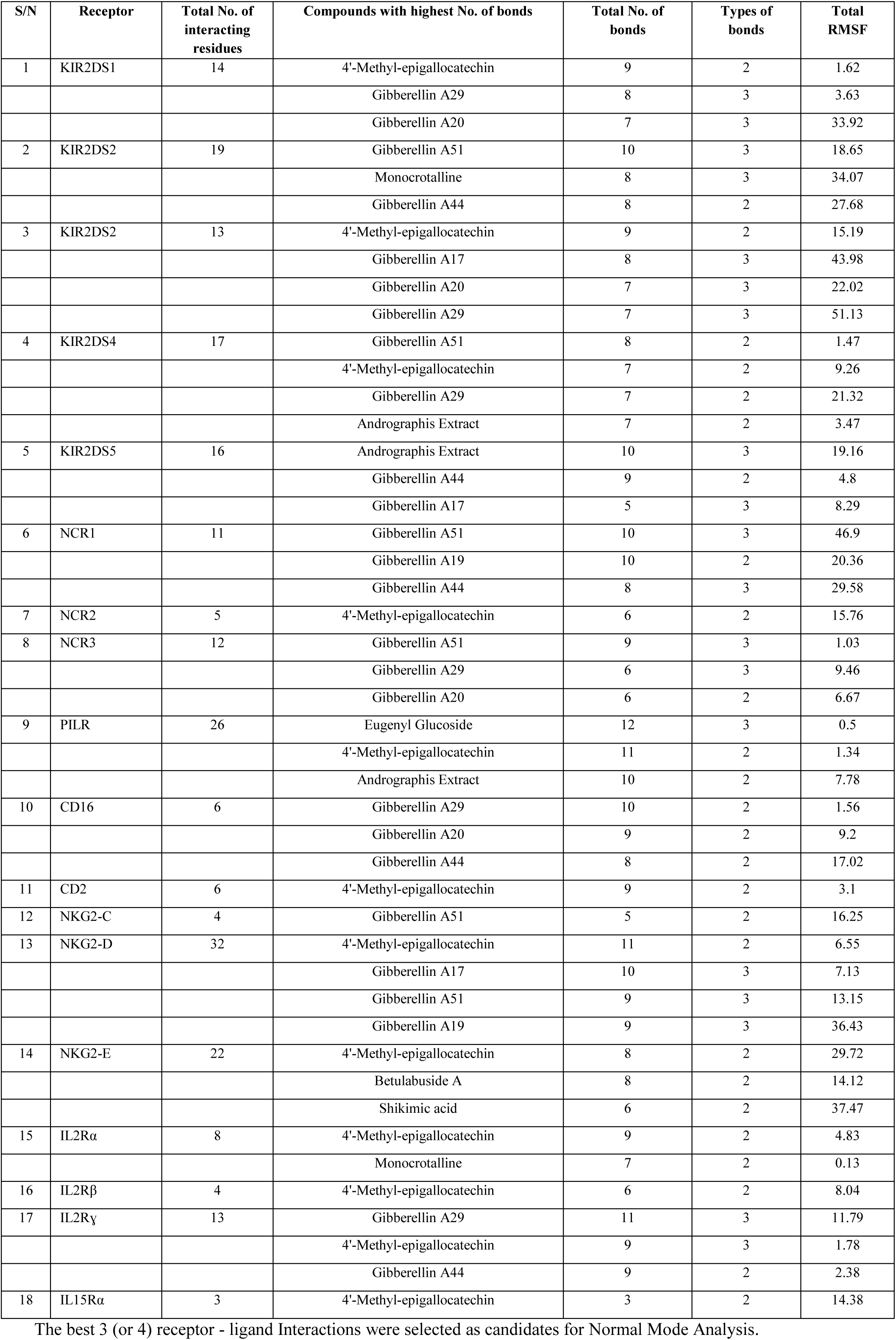
Summary of Receptor - Ligand Interactions and RMSF.

## 3.0 RESULTS AND DISCUSION

### 3.1 Preparation for docking

#### 3.1.1 Profiling and homology modeling of the protein structures

From Table 1, the four proteins modeled have very high percentage (between 85.96 and 92.44%) similarity with their templates. Usually, protein structures with over 30% identity to their templates can be predicted with an accuracy equivalent to a low-resolution X-ray structure [20]. In such high sequence identities, the major errors in modeling arise from the use of a poor template and inaccurate alignment of target-template sequence [21].

#### 3.1.2 Ramachandran Analysis

The ramanchandran plot was used to validate the macromolecular crystal structures of all the receptor targets to be studied by revealing the torsional conformation of their amino acids. From Table 2, all the protein structures have over 80% and 90% of their residues within the favoured and allowed regions respectively signifying good stereochemical quality. None of the proteins are intrinsically-disordered because of the chemical correctness of the torsional angles of their backbone [22].

When the φ and ψ angles are combined, an outlier residue has unusual torsional angles. All the protein structures had ramachandran outliers less than 0.05% signifying quality backbone conformation [23]. In this regard, the two proteins of least structural quality are KIR2DS1 and KIR2DS3 with 9 (0.046%) and 7 (0.036%) outliers respectively. These two proteins were homologically modeled from the same template, KIR3DL1. The relatively higher percentage of outliers found in these two proteins may be due to partially disordered large loops in the template. Loops have high electronic densities due to their structural flexibility and randomness and hence their residues show a broader range of dihedral angle values [24].

Though from Table 2, all the 18 proteins meet the required cut-off, IL15Rα and CD2 have the highest and lowest structural quality respectively. This difference is due to the method used for the structural analysis of these proteins. The structure of IL15Rα (pdb 4gs7) was obtained from x-ray crystallography, while CD2 (pdb1gya) was obtained from solution nuclear magnetic resonance (NMR). NMR gives a lesser resolving power than X-ray crystallography because it offers much more complex information from the same material. Most successful computational protein design use high-resolution X-ray crystallographic structures as templates [25].

#### 3.1.3 Energy minimization

As two non-bonding atoms in a protein structure approach, an atomic overlap (contact) occurs resulting in Van der Waals repulsion energy greater than 0.3 kcal/mol and subsequently leading to a steric clash. The webserver, *Chiron* is able to resolve severe steric clashes with minimal perturbation of the backbone of the native structure (less than 1 Å Cα RMSD).

*Chiron* generates a clash score which is a size-independent parameter obtained mathematically by the ratio of total VDW repulsion energy to the total number of contacts. From data generated from high-resolution structures, *Chiron* is able to determine if a protein has artifacts (excessive steric clashes) and return the clash score to physiological acceptability (0.02 kcal.mol−1.contact−1) [12].

A reduction in the total van der Waals (VDW) repulsion energy (Kcal/mol) of the clashing atoms. Would lead to a reduction in the steric clashes and consequently improve ligand-binding This is done computationally by rearranging this collection of non-bonding atoms in such a way that their inter-atomic forces are as close to zero as possible [12].

From Table 3, all 17 minimized structures have a physiologically acceptable clash ratio (clash score) of less than 0.02. There is no reduction in the total number of clashes and total VDW repulsion energy (Kcal/mol) in NCR1 and IL2Rγc signifying that these proteins already stable conformations. There is also no reduction in the steric clashes in all the protein structures that were modeled which are KIR2DS1, KIR2DS3, KIR2DS5 and NKG2E. This is because the *SWISS MODEL* webserver during the modeling process repairs distorted geometries or steric clashes through energy minimization [26]. IL2Rβ was not minimized probably due to missing heavy atoms of the backbone [12].

#### 3.1.4: Validation of docking protocol

1gya consists of CD2 and N-glycan (alpha-d-mannose, beta-d-mannose and N-acetyl-d-glucosamine) molecules. Figure 1 shows the images of the original 1gya and that of the separated, docked and superimposed. These two closely resemble thereby validating the docking protocol [27].

#### 3.1.5 Screening for Bioavailability

Prior to docking, a library of 1,697 compounds was screened for bioavailability using the Lipinski and Veber rules. The predictors of good oral bioavailability include number of rotatable bonds, hydrogen bond acceptors (≤ 10), hydrogen bond donors (≤ 5), molecular weight (≤ 500), low polar surface area (TPSA ≤ 140), and lipophilicity (Log P ≤ 5.0) [28, 29]. 1,048 front-runner compounds were selected with zero violations to both rules.

One limitation of the Lipinski rule is the fact that it only applies to compounds that are transported by diffusion through cell membranes. Actively transported compounds are exempted from this rule [30]. The conformational features of these compounds closely resemble endogenous metabolites and as such active transport is enhanced through ATP-dependent mechanisms [31]. This explains why so many proven compounds that have elicited *in vitro* cytotoxicity have been screened out [32].

### 3.2 Screening for potency

#### 3.2.1 Binding Affinities

For the purpose of screening, a uniform docking score of −7.0 kcal/mol was chosen as a cut-off value as this depicts strong protein-ligand binding. The choice of a higher docking score would increase the amount of data to be handled and also reduce potency [33]. The binding affinity values reveal the strength of ligand-protein interaction. After docking 1,048 ligands against 18 receptors, 377 front-runner compounds were selected as seen in Table 5 (summary of screening results). This implies that approximately 36% of the screened compounds obtained mainly from fruits, and vegetables have strong binding affinities with the activating receptors of the NK cells. This data further establishes the fact that phytochemicals of fruits, mushrooms and vegetables modulate NK cell activities and thereby promote the prevention of cancer [34].

Table 6 shows the summary of distributions and frequencies of receptor - ligand dockings at frequencies ≤-7.0 kcal/mol. NKG2D, NKG2E and PILR bound with the highest number of ligands in the library. NKG2D is known to be a promiscuous receptor and this suggests why it binds to a high number of ligands in the study [35]. NKG2E which was modeled with a NKG2A template (85.96% similarity) and NKG2D have similar hydrophobicity plots suggesting the possibility of promiscuity. PILR is also known to be a promiscuous type I transmembrane receptor and this suggests why it binds to a high number of ligands in the study [36].

On the contrary, Table 6 also reveals that NCR2, CD2, NKG2C, IL2Rβ and IL15Rα have less than 5%. This is suggestive of the fidelity of these receptors as they specifically bind to only a few ligands [37].

#### 3.2.2: Ligand Efficiency Metrics

LEM screening identifies compounds with greater potency and ADMET properties [38]. Maintaining the potency of a compound with the right molecular size and lipophilicity is a challenge in multi-parameter lead optimization. It is more ideal to optimize hits with the highest ligand efficiencies than those with the strongest binding affinities [39]. Table 5 reveals that a total of 192 front-runner compounds were obtained after screening using the LEM. The screened compounds had a LE of ≥0.3; an LELP of between −10 and 10; and LLE ≥ 5.0 [39]. Good LE values indicate that compounds have the desired potency at the appropriate weight. With lower molecular weight, there is also room for lead optimization to improve the potency and pharmacokinetic properties [40, 41]

### 3.3 QSAR-Based ADMET, Saturation and Promiscuity predictions

As seen in Table 5, a total of 69 front runner compounds emerged from the screening for saturation, promiscuity and pharmacokinetic properties. Many of the eliminated compounds remain viable candidates for lead optimization. Many of the eliminated compounds are also known to have strong antioxidant and immunomodulatory properties.

Molecular complexity which is measured by the carbon bond saturation (fraction of sp^3^ carbons - fsp^3^) plays a vital role drug discovery. Saturation directly correlates with solubility and saturated hydrocarbons have stability of the chemical bonds which makes them unreactive [42]. As seen in Table 8, all compounds with values less than 0.25 are unsaturated and therefore eliminated

While drug promiscuity may have its advantage, it elicits undesirable side effects due to ligand interactions with multiple protein targets in the biological system. A good predictor of promiscuity in bioassays is aggregation. Most drugs are not promiscuous even at high concentration. However, some have tendency to self-aggregate in aqueous media. These compounds have disruptive functional groups that can interfere with bioassays by causing activity artifacts leading to false positive results [43]. As seen in Table 8, there are no PAIN (Pan-assay Interference) compounds.

The absorption profile of a drug affects its bioavailability and consequently its efficacy and pharmacological effect [44]. Parameters such as water solubility, Caco-2 cell permeability, Human Intestinal Absorption (HIA), and Skin Permeability are within accepted range [45, 46, 47, 48]. Permeability glycoprotein (P-glycoprotein or Pgp) is a transporter protein that is located on the cell membrane. It is an ATP-dependent efflux pump which flushes out xenobiotics and toxic substances thereby limiting their cellular absorption [49]. From Table 7, all Pgp inhibitors were eliminated to avoid cellular toxicity. However, Pgp inhibitors can be used in overcoming multidrug resistance in cancers or administered with P-gp substrates to overcome the challenges of poor bioavailability associated with the later [50].

The Distribution of a drug determines the pharmacological effect and duration of action. From Table 7, the predicted distribution parameters such as steady state volume of distribution (VDss), Fraction unbound (Fu), Blood Brain Barrier (BBB) permeability and CNS permeability [51] are within pharmacological range

Many drugs that affect CYP450 enzymes by either inducing or inhibiting their activities. CY3A4 is the most abundant isoform in the liver. Inhibiting this enzyme can block it and cause an elevation of levels of substrate leading to toxicity or undesirable pharmacological effects [52, 53, 54]. From Table 8, all CYP450 enzyme inhibitors were eliminated.

The rate at which a drug is excreted determines the dose. Drug excretion is determined by such parameters as total Clearance (CL) which is a total of the renal clearance, hepatic clearance and the lung clearance. From Table 8, all lead compounds CL values within accepted pharmacological range. Human Organic Cation Transporter (OCT2) is a renal uptake transporter protein located on the proximal tubule cells. It removes mostly OCT2 substrates which are mostly cationic drugs from the blood into the urine. The concurrent administration of an OCT2 substrate with an OCT2 inhibitor would lead to a toxic intracellular accumulation of the OCT2 substrate. From Table 8, there is no OCT2 substrate

The toxicity profile of a drug is predicted based on QSAR models such as Microbial and fish toxicity, mutagenicity to *Salmonella typhimurium* (Ames Test), Human ether-a-go-go-related gene (hERG) inhibition, Skin Sensitization, Hepatotoxicity. All lead compounds were non mutagens, non-hERG inhibitors and non-dermatoxic. From Table 9, Eugenyl Glucoside, Gibberellin A19, Gibberellin A51 and Gibberellin A53 are predicted to be hepatotoxic. This implies that they possess structural moieties that could elicit the disruption of normal liver function. This kind of hepatotoxicity usually has a predictable dose-response curve. This suggests that doses below the MTD cannot induce hepatotoxicity [55]. Other dose related toxicity indicators which include Microbial and fish toxicity, Maximum Tolerated Dose (MTD), Acute Toxicity (LD50), and Chronic Toxicity are within acceptable pharmacological range.

### 3.4 Bioactivity

Affinity does not necessarily predict activity. Binding ligands could be either agonists or competitive inhibitors. Based on a particular drug target, a compound is considered to be active when it’s a bioactivity score is more than 0.0; moderately active when score is between −5.0 and 0.0; and inactive when the score is less than −5.0 [56].

Table 10 reveals 17 compounds that are active as nuclear receptor ligands. Many of these compounds are multi-targeted, binding to multiple receptor targets. Bioactivity screening also eliminates promiscuous binding compounds as seen in PILR, NKG2E and NKG2D receptors.

### 3.5 Specificity-Promiscuity Analyses

There is no correlation between potency and specificity. Selectivity plays a strategic role in drug development [57]. Beyond potency, the selectivity of a drug is also important as this guarantees specificity at the biological target reducing unwanted side effects [58].

From Table 11, the comparative analysis of binding affinities shows 6 compounds that have absolute binding specificity with a single receptor (NKG2D or NKG2E) at ≤ −7.0 kcal/mol. Specificity also depicts the strength of interaction between ligand and protein. High chemical specificity means that proteins bind to a limited number of ligands. This is important as certain physiological processes might require specificity [59, 60]. Compounds such as 4’-Methyl-epigallocatechin, Andrographis Extract, and Gibberellins A17, 19, 29 & 44 have strong binding affinities with 15 and above receptor protein targets.

### 3.6 Similarity analysis

Structural similarity may suggest closeness in biological activity [61].

#### 3.6.1 Ligand Similarity analyses

As seen in Table 12, a pairwise ligand similarity analyses of Gibberellins A17, A19, A20, A29, A44, A51 and A53 reveal Tanimoto scores ranging from 0.50 to 0.81. Gibberellins A20 and A51 have the same chemical formula but different stereochemistry. Carveol & cis-Carvotanacetol have a Tanimoto score of 0.51. These compounds have been predicted to elicit similar function and would be useful in building pharmacophores for ligand-based drug design [62].

#### 3.6.2: Receptor similarity

The multi-target binding of the ligands is likely due to the structural similarities of the protein targets. Empirical evidences show that ligands could have the same binding pocket in different proteins [57]. This may be due to genetic similarities of the proteins. Isoforms of the same protein and those that by co-evolution may exhibit similar biochemical reactions might have the same binding sites [63].

The structural similarity of the target protein was studied using a percent identity matrix in Table 13. Amino acid sequence alignments that produce a pairwise sequence identity >40% is considered high [64]. Out of the 11 members of the Immunoglobulin super family of receptors, KIR2DS1, KIR2DS2, KIR2DS3, KIR2DS4, and KIR2DS5 are highly similar proteins as they have degree of conservation ranging from 86.53-92.65% (Table 13). Of all the 3 lectin-like receptors, consensus sequences only exist between NKG2E & NKG2C with a 90.04% identity. This signifies that these two sets of proteins are isoforms. NKG2E and CD2 have the least identity of 6.36%.

From Figure 2, all the receptors have a common ancestor and have evolutionary relatedness. An original speciation event occurred resulting in three lineages (roots). The tree also depicts the direction of evolution, with the flow of genetic information moving from the roots, through the clades, to the branches, to the taxa and outgroups. Root 3 consists exclusively of the KIR2DS series of receptors. Root 1 Clade 2 also consists of all the lectin-like receptors. Most closely related pairs exist in the sister taxa. KIR2DS1-5 are the most closely related family in all the 18 receptors. The NCR 1-3 are the most divergent.

### 3.7 Binding Site analyses

All residues that are involved in the ligand–protein interactions are located within the extracellular domains of the receptor (See supplementary data). Receptor signaling should commence from the extracellular domain through the helical domain to the cytoplasmic domain.

The greater the number of ligand interactions within the functional domains, the greater the biological activity of the protein is triggered. The 18 receptor targets have functional domains such as Immunoglobulin-like (C and V types), sushi, C-type lectin, and fibronectin type III domains. For example, as seen in supplementary data, Gibberellin A53, 4’-Methyl-epigallocatechin and Gibberellin A51 have all their interactions (hydrophobic and hydrogen bonds) within the C2 type 1 and C2 type 2 domains of the KIR2DS4 receptor (N.B. A value of 5 should be added to all the residue numbers for KIR2DS4 to take care of the rearrangement during energy minimization.)

IL2Rβ has 5 binding sites. Proteins with multiple binding sites show cooperativity. The assembly of the IL2R-IL15R complex allows interfaces between these proteins to create hydrophobic pockets for ligand binding. However, the binding at the original site affects the affinity of all the other sites [65].

### 3.8: Normal Mode Analysis

Protein flexibility is determined by fluctuations of the alpha carbon atoms of the amino acids. This is seen as rearrangements of side chains or changes in the backbone. Ligand binding induces conformational changes in the protein structure [66]. The stability of protein-ligand complexes would impact on protein function. As revealed in Table 14 structures with the lowest global fluctuation are indicative of the most stable protein-ligand complexes. Ligands of these most stable complexes are the most suitable drug candidates for their respective receptors. The highest numbers of interacting residues are seen in NKG2-D, PILR and NKG2-E which have 32, 26 and 22 residues respectively. The lowest RMSF value is seen in the interaction between IL2Rα and Monocrotalline (0.13), while the highest is between KIR2DS2 and Gibberellin A29 (51.13). The highest number of bonds is seen between PILR and Eugenyl Glucoside (12).

### 3.9 Brief review of successful leads

#### 3.9.1: Andrographis Extract

obtained from *Andrographis Paniculata* (King of Bitters) has exhibited potent anti-inflammatory and anticancer properties. Its chemo-preventive activity is revealed in the growth suppression of cancer cells by inducing apoptosis and by inhibiting PI3K/AKT, NF-kappa B, and other kinase pathways [67].

In mice, the ethanol extract of *Andrographis paniculata* also significantly induced antibody production and delayed type hypersensitivity response to sheep red blood cells. In terms of nonspecific immune response, the Andrographis extract induced significant immunostimulation as measured by proliferation of splenic lymphocytes, thymocytes and bone marrow cells; the migration of macrophages and phagocytic activity [68,69].

*Andrographis paniculata* extract is known to be one of the natural products that enhance the efficiency of NK cells in the control of cancer. It promotes NK cell mediated lysis of metastatic tumor cells in mice through an antibody-dependent complement-mediated cytotoxicity [69, 70, 71]. It also significantly increases the production of interleukin-2 and interferon-gamma and decreases pro-inflammatory cytokines such as TNF-α, GM-CSF, IL-1ß, and IL-6 in tumour-bearing animals [69, 70].

#### 3.9.2 The Gibberellins A17, A19, A20, A29, A44, A51 & A53

Gibberellins (GAs) are a group of closely related plant hormones that regulate several physiological and developmental processes which include germination, elongation, flowering and fruiting [72]. Gibberellins can be obtained from *Abelmoschus esculentus* (Okro) and *Pisum sativum* (Green peas) [73,74].

Gibberellin has been implicated as a modulator of the plant innate immunity. It plays significant role in plant-microbe interaction especially as it has to do with the root’s basal defense. Successful fungal colonization is due to altering gibberellin signaling in plants [75]. Gibberellin modulates plant immune system by regulating the Salicylic acid (SA), Jasmonic acid (JA) and Ethylene (ET) signaling systems [76].

There were no direct cytotoxic effects of Gibberellins A17, A19, A20, A29, A44, A51 & A53 found in literature. However, Gibberellin derivatives such as GA-13315 reveal strong antineoplastic effects both *in vitro* and *in vivo*. It inhibits the growth and also accelerates the apoptosis of KB oral cancer cells. GA-13315 also possesses anti-angiogenic properties [77, 78].

GA-13315 inhibits the P-glycoprotein thereby reducing multidrug resistance induced by cancer cells and it also triggers the multidrug resistance-associated Protein −1 [79]. Other synthesized gibberellin derivatives bearing two alpha, beta-unsaturated ketone units showed strong activity in MTT assay against A549, HepG2, HT29, and MKN28 human cancer cell lines. They also exhibited inhibition to topoisomerase I activity [80].

Gibberellin A4 is known to be a native ligand to the Fab fragment of the haptenic mouse monoclonal antibody, 4-B8 (8)/E9. X ray crystallography of the Fab fragment reveals a typical beta barrel fold which is a common motif of all immunoglobulins [81]. This suggests why Gibberellins might be able to bind to Immunoglobulin-like receptors which have immunoglobulin domains.

#### 3.9.3–4’-Methyl-epigallocatechin

This compound can be found in Locust beans (*Parkia biglobosa*). Epigallocatechin which is found in Green Tea *Camellia sinensis* can also be methylated into 4’-Methyl-epigallocatechin in the human body [82, 83].

Another epigallocatechin derivative such as epigallocatechin gallate (EGCG) which is also found in Green Tea has anticancer effects. Through cell mediated immunity, EGCG reverses myeloid-derived suppressor cell activity [84,85]. ECCG is also able to modulate both the innate and adaptive immune systems. In ameliorating experimental arthritis in mice, it upregulates the Nrf-2 antioxidant pathway, induces Indoleamine-2, 3-dioxygenase (IDO)-producing dendritic cells and increases Treg population [86].

#### 3.9.4 Shikimic Acid

is a cyclohexanecarboxylic acid, obtained from *Malus domestica*, Apples. It exhibits anti-inflammatory and antioxidant activities [87,88]. Shikimic acid complex of platinum (II) is active against Leukemia (L1210 and P388) and B 16 Melanoma cell lines [89]. The shikimic acid-based synthesis of zeylenone is widely used as a preparation for chemotherapy in cancer patients [90]. Shikimic acid analogue skeleton is a constituent of several antitumor products [91].

## 4.0 CONCLUSION

With the aim of triggering cytotoxicity, 1,697 natural compounds derived from 83 plants were docked against 18 activating NKC receptor targets. After rigorous screening, 17 bioactive, non-promiscuous hit compounds with good physicochemical and pharmacokinetic properties were identified.

To add value to the drug discovery process, lead optimization may be necessary in order to adjust the structures of the compounds to achieve stronger binding affinity, greater potency and better ADMET-prediction. The identification of the pharmacophores of strong binding affinity-compounds and the modification of their core structural moieties, could achieve the ideal pharmacokinetic properties.

A further molecular dynamics simulation study is required to confirm the viability of the 18 drug targets. With the right parameterization, the strength and sustainability of the molecular interactions between these proteins and the lead compounds.

